# A study of Kibbutzim in Israel reveals risk factors for cardiometabolic traits and subtle population structure

**DOI:** 10.1101/239509

**Authors:** Einat Granot-Hershkovitz, David Karasik, Yechiel Friedlander, Laura Rodriguez-Murillo, Rajkumar Dorajoo, Jianjun Liu, Anshuman Sewda, Inga Peter, Shai Carmi, Hagit Hochner

## Abstract

Genetic studies in isolated populations have provided increased power for identifying loci associated with complex diseases and traits. We present here the Kibbutzim Family Study (KFS), initiated for investigating environmental and genetic determinants of cardiometabolic traits in extended Israeli families living in communes characterized by long-term social stability and homogeneous environment. Extensive information on cardiometabolic traits, as well as genome-wide genetic data, was collected on 901 individuals, making this study, to the best of our knowledge, the largest of its kind in Israel. We have thoroughly characterized the KFS genetic structure, observing that most participants were of Ashkenazi Jewish (AJ) origin, and confirming a recent severe bottleneck in their recent history (point estimates: effective size ≈450 individuals, 23 generations ago). Focusing on genetic variants enriched in KFS compared with non-Finnish Europeans, we demonstrated that AJ-specific variants are largely involved in cancer-related pathways. Using linear mixed models, we conducted an association study of these enriched variants with 16 cardiometabolic traits. We found 24 variants to be significantly associated with cardiometabolic traits. The strongest association, which we also replicated, was between a variant upstream of the *MSRA* gene, ≈200-fold enriched in KFS, and weight (P=3.6·10^−8^). In summary, the KFS is a valuable resource for the study of the population genetics of Israel as well as the genetics of cardiometabolic traits in a homogeneous environment.

## Introduction

From an evolutionary perspective, rare variants are on average more recent than common variants, and are thus more likely to be disease risk factors. Genetic association studies of common diseases and traits in isolated populations were particularly advantageous for identification of risk loci that are rare in the general population, but possibly enriched or common in an isolated population (Trecartin *et al*. 1981; Baier and L.Hanson 2004; Kristiansson *et al*. 2008; Sabatti *et al*. 2009; Zeggini 2014; Nair and Baier 2015; Fang *et al*. 2016; Gilly *et al*. 2016; Lopes *et al*. 2016; Zeggini *et al*. 2016). The Ashkenazi Jewish (AJ) population has been an attractive population for genetic studies, because of its unique demographic history of a recent severe bottleneck followed by a rapid expansion and monogamy (Guha *et al*. 2012). AJ were found to carry unique mutations responsible for a wide range of Mendelian disorders as well as risk factors for complex diseases (Charrow 2004; Kenny *et al*. 2012). Importantly, while these mutations may be unique or nearly-unique to AJ, they often highlight pathways of broad significance.

Cardiovascular disease (CVD) is a common cause of death worldwide (Mathers and Loncar 2006). Genome-wide association studies (GWAS) in unrelated individuals have identified thousands of genetic variants associated with CVD and numerous related risk factors (Fall and Ingelsson 2014; Atanasovska *et al*. 2015), but the risk is not fully explained. The Kibbutzim Family Study (KFS) was established in 1992 to investigate the environmental and genetic basis of cardiometabolic risk factors and their change over time (Friedlander *et al*. 1995, 2005, 2006; Lemaitre *et al*. 2008). The participants belonged, at the time of recruitment, to large families living in close-knit communities, called “Kibbutzim”, in Northern Israel. The Kibbutz has been a communal settlement, which has created a relatively homogeneous environment for its members. For example, earnings were uniformly distributed, and Kibbutz members typically dined jointly. Kibbutz members are mostly of Ashkenazi Jewish ancestry, with the remaining members belonging to other Jewish subgroups. The KFS is thus expected to be a useful resource for the study of cardiometabolic genetic risk factors.

While most association studies so far were conducted on unrelated individuals, the extended family design of the KFS has the advantages of reduced sensitivity to population stratification bias and the ability to detect Mendelian inconsistencies (Ott *et al*. 2011). Also, as in isolated populations, family-based studies have additional ability to enrich for genetic loci containing rare variants. Familial aggregation, segregation analyses and linkage and candidate gene association studies were previously conducted in KFS (Friedlander *et al*. 1999a; b, 2003, 2005, 2006; Sinnreich *et al*. 1999; Lemaitre *et al*. 2008), focusing on outcomes such as LDL peak particle diameter (Friedlander *et al*. 1999a), fibrinogen variability (Friedlander *et al*. 2003) and red blood cell membrane fatty acid composition (Lemaitre *et al*. 2008).

Here, we present results for genome-wide genotyping of 901 KFS participants (genotyped at ≈500k variants). We aimed to (1) characterize the population genetics of the KFS population and its relation to other worldwide populations; and (2) assess the contribution of enriched genetic variants in the KFS to the genetic basis of cardiometabolic traits and other health-related phenotypes.

## Methods

### Recruitment strategy

Participants (n=1033) in the KFS were recruited and sampled in two phases in 1992–1993 and 1999–2000 (Friedlander *et al*. 1999a, 2006), from six Kibbutzim in Northern Israel (Friedlander *et al*. 2003). The first recruitment phase of the study (1992–1993) included 80 extended families, ranging in size from 2 to 43 individuals (Sinnreich *et al*. 1998). During the second phase (1999–2000), participants from the first phase were all invited for repeat examinations (80% response rate) and new participants were recruited, giving a total of 150 extended families ranging in size from 2 to 55 individuals (Friedlander *et al*. 2006). Families were invited to participate if they consisted of at least four individuals who (i) lived in the Kibbutz, (ii) spanned at least two generations, and (iii) were at least 15 years old. Families were retained if at least two family members consented to participate in the study. Overall, 1033 participants were recruited; 111 were examined only in the first phase, 533 only in the second phase, and 389 were included in both (Figure 1).

**Figure 1:**
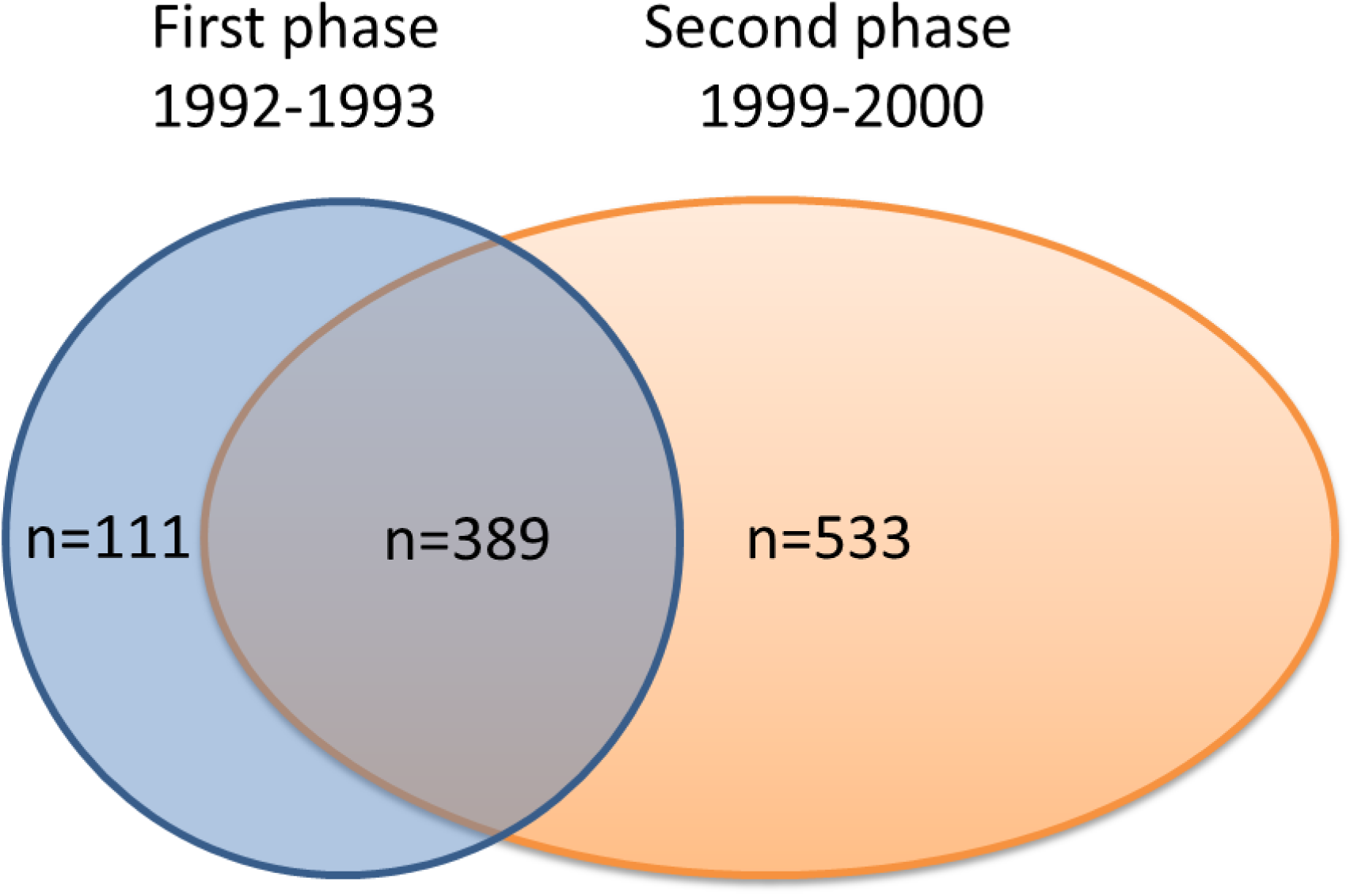
The Kibbutzim Family Study sample sizes at each phase.

### Data Collection

All subjects signed an informed consent and completed a self-administered socio-demographic and health questionnaire, including questions on medical and family history, cigarette smoking, alcohol consumption, and physical activity (Sinnreich *et al*. 1998; Friedlander *et al*. 2006). The participants also signed a set of psychosocial questionnaires, similar to that described by (Kark *et al*. 1996), with some differences between the phases. Dietary information was collected using a Food Frequency Questionnaire in the second phase only. Peripheral blood samples (25 ml) were collected at both phases following a 12-hour fast. Anthropometric and blood pressure traits (described below) were measured at both phases (Friedlander *et al*. 1995; Lemaitre *et al*. 2008).

### Genotyping and quality control

Of the 1033 participants recruited, 938 had high quality DNA samples (A260/280 >1.8, concentration >50ng/μl). Genotyping was performed using Illumina HumanCoreExome BeadChip, consisting of ≈240,000 tag single nucleotide polymorphisms (SNPs) and ≈240,000 exome variants, including loss of function variants and indels (https://support.illumina.com/array/array_kits/humancore_exome_beadchip_kit.html).

Standard quality control (QC) procedures were applied to filter variants and individuals using Plink 1.90 (https://www.cog-genomics.org/plink2) (Chang *et al*. 2015). We removed variants with genotyping rate <90%, Mendelian error rate >10%, and significant deviation from Hardy–Weinberg equilibrium (P<10^−6^). We removed individual samples with genotyping rate <95%, unresolved gender mismatch, autosomal heterozygosity >5%, and Mendelian error rate >5%. We confirmed the reported familial relationships using identical-by-descent (IBD) segments (see below). The concordance across 22 DNA duplicates was 99.93%-99.96%, and the sample with the lower call rate for each pair was excluded. We calculated runs of homozygosity (ROH) using plink (--homozyg-kb 5000). A total of 901 individuals and 323,708 variants (281,586 variants with minor allele frequency (MAF) > 1%) passed QC and were used in downstream analyses.

### Principal Component Analysis

Principal component analysis (PCA) was performed using the PC-AiR (Principal Components Analysis in Related Samples) method in the R GENESIS package (Conomos *et al*. 2016), which is robust to known or cryptic relatedness. Our reference panel included West-Eurasian populations (covering Europe, the Middle East, and the Caucasus, n=922) (Behar *et al*. 2013) and the Jewish groups, listed in Supplementary Table 1 (n=174) (Behar *et al*. 2013). These samples served as the “unrelated subset” for running PC-AiR, and principal components (PC) values were then computed for the KFS individuals (the “related subset”). We also performed another analysis using only the Jewish groups (n=174) as the reference population, in order to focus on substructure within the Jewish ancestries. In that analysis, we also included a panel of AJ recruited in the United States for The Ashkenazi Genome Consortium (TAGC) (n=128) (Carmi *et al*. 2014), which allowed us to examine differences in ancestry between AJ from Israel (KFS) and the United States. Another analysis was performed using only the AJ samples from Behar et al., 2013 (n=29) as the reference population, to focus on differences between Western and Eastern AJ.

Variants used in the PC-AiR analysis were restricted to MAF>1% and were pruned to eliminate linkage disequilibrium (LD), using the --indep-pairwise command in Plink (window size 50kb, variant count of 10 for shifting the window at the end of each step, and LD between variants (*r*^2^) <0.1).

### IBD Sharing and demographic reconstruction

The genotypes were phased using SHAPEIT2 v2 (O’Connell *et al*. 2014). When SHAPEIT2 is run on family data, explicit family information is first ignored, and then a hidden Markov model (HMM) is applied (duoHMM) to correct switch errors in the inferred haplotypes according to parent-child relationships (O’Connell *et al*. 2014). We detected pairwise shared haplotypes (IBD segments) in the cohort using GERMLINE (Gusev *et al*. 2009). The GERMLINE parameters were: bits=40, err_hom=1, err_het=1, min_m=3 (centiMorgan, cM), hextend=0. The segments identified by GERMLINE are known to suffer from a high rate of false positives (Durand *et al*. 2014). We thus filtered the segments using HaploScore (Durand *et al*. 2014), which is a measure of the number of genotyping and phase switch errors required to explain each segment; segments with a HaploScore >2.5 were removed. We also removed segments with more than 5% overlap with sequence gaps (http://genome.ucsc.edu/), less than 16 variants per cM, or an overlap with the HLA region (chr6:24-37M). For the demographic analysis in the AJ samples, we retained only segments shared between Ashkenazi founders (n=303), as identified by the first PC in the PCA of Jewish populations (Results). An evaluation of the improvement in the accuracy of detected IBD segments due to the pedigree-based phasing is provided in Supplementary Note 1.

To estimate the demographic history of AJ using the IBD segments, we used a previously developed method (Palamara *et al*. 2012), which we have recently applied (Carmi *et al*. 2014; Zidan *et al*. 2015; Gilbert *et al*. 2017). Briefly, we assumed a model of an ancestral (diploid) effective population size N_A_, then an instantaneous bottleneck, T_B_ generations ago, with an effective population size N_B_, and finally an exponential expansion to a current population size N_C_ (Zidan *et al*. 2015). We divided the space of segment lengths between 3-30 cM into 25 intervals equally spaced in log-scale. For each interval, we computed the proportion of the genome, averaged over all each pairs of haplotypes, that falls within IBD segments with length in the specified interval. We used a grid search to infer the demographic parameters. For each parameter combination and for each interval, we computed the expected proportion of the genome in IBD segments using a previously developed theory (Palamara *et al*. 2012) (see also (Zidan *et al*. 2015)). The fit of each model was evaluated as the sum, over all intervals, of the square of the log-ratio of the actual (AJ) and theoretical proportions. The fitting error was plotted for each parameter separately, by fixing the value of the given parameter and optimizing the model over the other three parameters.

### Imputation

After pre-phasing the microarray genotyped data using SHAPEIT2, imputation was performed in chunks of length 5MB each, using IMPUTE2 (Howie *et al*. 2012). For the reference panel, we used either an Ashkenazi-only reference panel (TAGC; n=128) (Carmi *et al*. 2014), the 1000 Genomes reference panel phase 1 version 3 (n=1092), or the combined Ashkenazi + 1000 Genomes panel (n=1220). We ensured strand consistency of the array genotypes compared to the sequence data. To compare the imputation accuracy between the three reference panels, we used the estimates provided by IMPUTE2, as the concordance between true array genotypes and their imputed value when masked. The concordance was highest when the imputation was performed using the combined reference panel. Therefore, we used the combined panel for our association analyses. Imputed genotypes were available at 82,328,870 variants; however, for most analyses we only considered 6,858,900 variants with MAF ≥ 1% (calculated in founders only) and imputation quality score ≥0.9.

### Enrichment Analysis and Functional Annotation

In populations that are genetically isolated or have undergone recent strong genetic drift such as Ashkenazi Jews (Palamara *et al*. 2012; Carmi *et al*. 2014), it is expected that some risk variants of large effects may rise in frequency compared to the general population, resulting in an increased power to detect an association (Hatzikotoulas *et al*. 2014). We thus focused on variants with a substantially higher frequency in the founder population (in our case, the KFS participants) compared to the general population, which we take as the non-Finnish Europeans (NFE) population from The Genome Aggregation Database (gnomAD). We observed that a naíve search for variants with a large frequency difference between the KFS and Europeans led to numerous artifacts. We thus implemented a series of QC steps. First, we filtered variants for which the KFS MAF (founders only, n=393) differed by >10% from the AJ MAF in gnomAD (Lek *et al*. 2016). Similarly, we filtered variants with >10% MAF difference between the NFE population from the 1000 Genomes Project phase 3 (CEU + GBR + TSI + IBS; n=404) (Auton *et al*. 2015) and gnomAD (n≈7500) (Lek *et al*. 2016). Lastly, we focused on variants that are very rare (MAF<0.1%) in the gnomAD NFE population and relatively common (MAF≥1%) in the KFS cohort, resulting in n= 212,505 “enriched” variants.

To determine if the MAF ratio correlates with functional consequences of the enriched variants, we annotated them using the SnpEff version 4.3q (Cingolani *et al*. 2012). Variants that were present in the KFS but missing in the gnomAD NFE population (i.e. “infinite-fold” enriched) were excluded from the functional annotation. On a subset of variants with >100x KFS/NFE frequency ratio and high/moderate predicted functional impact, we also performed gene-set enrichment analysis (GSEA) using the Molecular Signatures Database (MSigDB, Subramanian et al., 2005).

### Association analysis

Association tests were carried out in BOLT-LMM v2.2 (Loh *et al*. 2015b). BOLT-LMM uses a linear mixed model approach, which accounts for relationships between individuals and population structure, as well as handles the imputed ‘dosage’ data. For building the mixed model, we used 299,509 genotyped variants (MAF>0.1%). We used the 1000 Genomes LD-Score table provided with the BOLT-LMM software package. Of the 6,858,900 imputed genetic variants available after QC, we focused on 212,505 “enriched” variants with 10x higher MAF in KFS compared to Europeans (see above).

To calculate the P-value threshold for genome-wide significance, we observed that most of the enriched variants were clustered in long haplotypes. Indeed, after LD pruning in Plink (*r*^2^>0.2), only 16.9% of the enriched variants remained, suggesting that the enriched variants are densely distributed in specific haplotypes, as opposed to sparsely genome-wide. We thus proceeded as if the enriched variants cover a contiguous proportion of 212,505/6,858,900 = 0.031 of the genome. The conventional genome-wide threshold, P<5·10^−8^, is based on an effective number of ≈1 million effective tests genome-wide (Pe’er *et al*. 2008). Thus, our enriched variants represent approximately 10^6^·0.031 = 3.1·10^4^ tests. This leads to a suggestive genome-wide significance threshold of 0.05/3.1·10^4^ = 1.61·10^−6^.

Supplementary Table 3 lists the 16 anthropometric and cardiometabolic traits we analyzed and their corresponding heritability estimates (based on all imputed genetic variants), as calculated by BOLT-REML (Loh *et al*. 2015a). In case a phenotype was measured in both phases, we used the value from the most recent phase. All association models used age, gender, and phase as covariates. Lipid-lowering medication was accounted for in analyses of lipid traits by introducing a dichotomous covariate for medication use (yes/no). We adjusted for blood-pressure-lowering medication by adding 10 and 5 mmHg to systolic (SBP) and diastolic (DBP) blood pressures, respectively (Cui *et al*. 2003). Lipoprotein (a), C-reactive protein, and triglycerides variables were inverse normal transformed because of their skewed distributions. We observed no improvement in association results when using the non-infinitesimal mixed model test in BOLT-LMM, and thus all reported results are for the standard infinitesimal model.

## Results

### Samples

Our study included 1033 participants (47% male, 53% female), recruited during two phases (1992–1993 and 1999–2000) from 150 families (445 founders; 43%); see Methods for details. The mean number of generations per family was 2.63 (range 1–4), with the majority of families spanning 3 (57.3%) and 2 (28.0%) generations. The mean family size was 6.89 individuals (range 2–55).

Participants’ characteristics by gender are given in Table 1. The mean age was 44±20 yrs (similarly between men and women) and 73% were born in Israel. About half the individuals attained >12 years of schooling, in both males and females, and about 10% reported being current smokers. The majority of individuals reported no engagement in strenuous physical activity at leisure time, and similarly for physical activity during work. The 16 anthropometric and cardiometabolic traits we used for the association analysis are summarized in Supplementary Table 4 by gender and age group (≤29 years, 29–59 years, and ≥59 years).

**Table 1:**
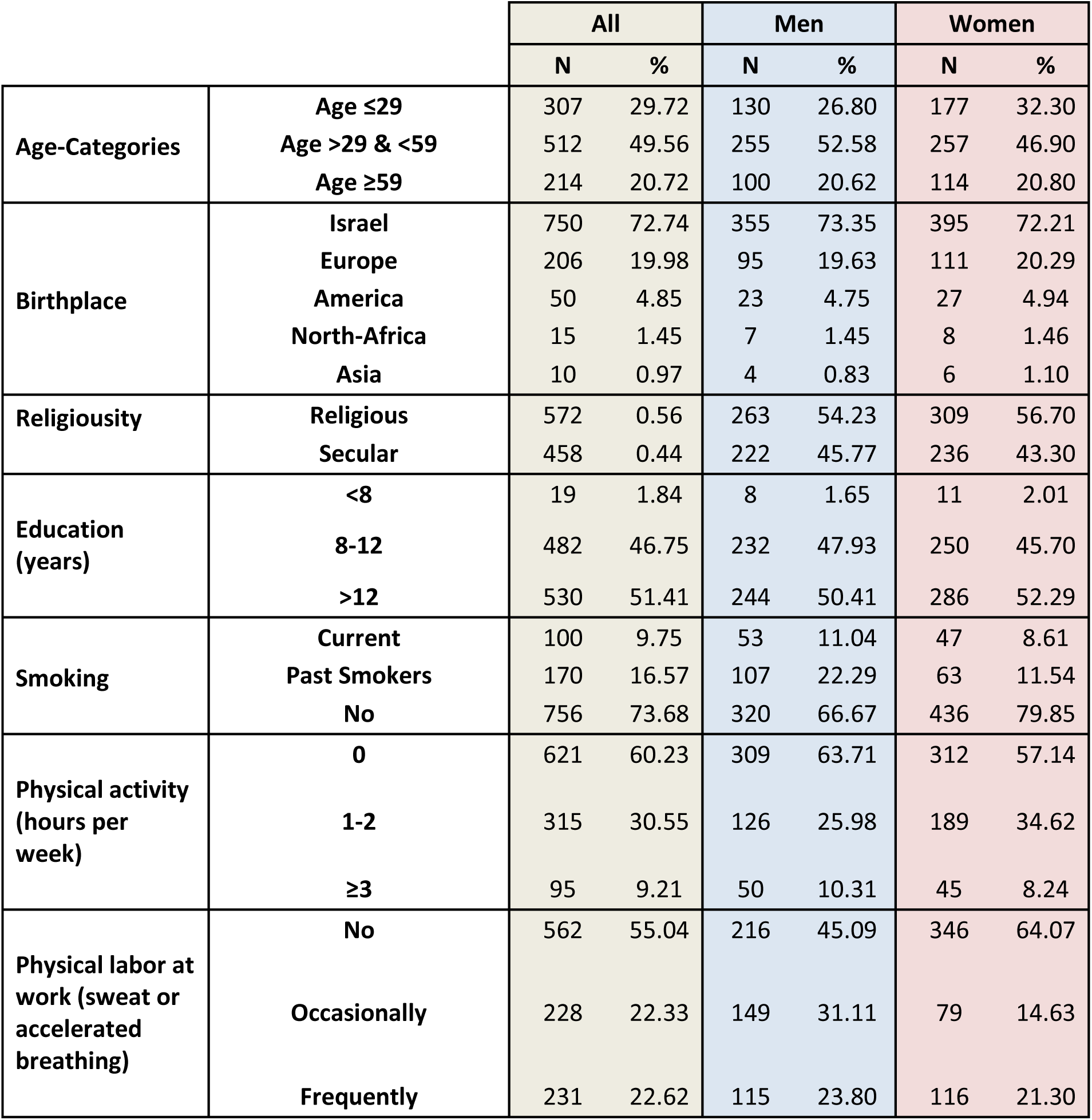
Socio-demographic characteristics of the KFS.

|  |  | All |  | Men |  | Women |  |
| --- | --- | --- | --- | --- | --- | --- | --- |
|  |  | N | % | N | % | N | % |
| Age-Categories | Age ≤29 | 307 | 29.72 | 130 | 26.80 | 177 | 32.30 |
|  | Age >29 & <59 | 512 | 49.56 | 255 | 52.58 | 257 | 46.90 |
|  | Age ≥59 | 214 | 20.72 | 100 | 20.62 | 114 | 20.80 |
| Birthplace | Israel | 750 | 72.74 | 355 | 73.35 | 395 | 72.21 |
|  | Europe | 206 | 19.98 | 95 | 19.63 | 111 | 20.29 |
|  | America | 50 | 4.85 | 23 | 4.75 | 27 | 4.94 |
|  | North-Africa | 15 | 1.45 | 7 | 1.45 | 8 | 1.46 |
|  | Asia | 10 | 0.97 | 4 | 0.83 | 6 | 1.10 |
| Religiosity | Religious | 572 | 0.56 | 263 | 54.23 | 309 | 56.70 |
|  | Secular | 458 | 0.44 | 222 | 45.77 | 236 | 43.30 |
| Education (years) | <8 | 19 | 1.84 | 8 | 1.65 | 11 | 2.01 |
|  | 8-12 | 482 | 46.75 | 232 | 47.93 | 250 | 45.70 |
|  | >12 | 530 | 51.41 | 244 | 50.41 | 286 | 52.29 |
| Smoking | Current | 100 | 9.75 | 53 | 11.04 | 47 | 8.61 |
|  | Past Smokers | 170 | 16.57 | 107 | 22.29 | 63 | 11.54 |
|  | No | 756 | 73.68 | 320 | 66.67 | 436 | 79.85 |
| Physical activity (hours per week) | 0 | 621 | 60.23 | 309 | 63.71 | 312 | 57.14 |
|  | 1-2 | 315 | 30.55 | 126 | 25.98 | 189 | 34.62 |
|  | ≥3 | 95 | 9.21 | 50 | 10.31 | 45 | 8.24 |
| Physical labor at work (sweat or accelerated breathing) | No | 562 | 55.04 | 216 | 45.09 | 346 | 64.07 |
|  | Occasionally | 228 | 22.33 | 149 | 31.11 | 79 | 14.63 |
|  | Frequently | 231 | 22.62 | 115 | 23.80 | 116 | 21.30 |

We genotyped 938 individuals using the Illumina HumanCoreExome BeadChip. After quality control, 901 individuals and 323,708 variants (281,586 variants with minor allele frequency (MAF) > 1%) remained and were used in the population genetics analyses (Methods). After imputation with IMPUTE2 and a combined Ashkenazi Jewish and cosmopolitan reference panel, 6,858,900 variants with MAF ≥ 1% were available for the association analyses (Methods).

### Population genetics

#### Principal Component Analysis (PCA)

To study the genetic ancestry of the KFS participants, we ran PCA (Methods) on the genotyped KFS samples (n=901), along with worldwide (n=922) and Jewish (n=174) reference populations (Behar *et al*. 2013) (Supplementary Table 1). The first two principal components (Figure 2) distinguish three main non-Jewish population groups: European, Caucasian, and Middle-Eastern, and six Jewish populations: Ashkenazi, Sephardi, North-African, Yemenite, Middle-Eastern, and Caucasian. A partial overlap is observed between AJ and European non-Jews, as well between Middle-Eastern and Caucasian Jewish and non-Jewish populations. The KFS samples largely overlap with the AJ reference samples (Behar *et al*. 2013). To study the non-Ashkenazi ancestries in the KFS, we ran PCA with the KFS samples and the Jewish reference populations only (Figure 3). The number of individuals with exclusive AJ ancestry, as distinguished by the first PC (PC1 ≤ 0), was n=733 (81.4%). The majority of the remaining individuals overlapped with the Sephardi and North-African Jewish clusters, but the Middle-Eastern, Caucasian, and Yemenite Jewish populations were also represented. Some individuals seemed to have mixed Ashkenazi and other Jewish ancestry, although quantifying their exact number is difficult with PCA. Self-reported country of birth allowed us to compare the PCA-based and self-reported Jewish ancestry for 247 individuals born outside of Israel (Supplementary Figure 1). Among 140 individuals self-reported as AJ (born in Austria, Belgium, Denmark, England, Germany, Holland, Hungary, Ireland, Poland, Romania, Russia, Sweden, or Switzerland), 136 (97%) met the defined genetic criterion (PC1 ≤ 0). Among the 11 individuals self-reported as North-African Jewish (born in Tunisia, Morocco, Algeria, and Libya), 9 (82%) met a pre-defined genetic criterion (PC1>0.03 and PC2>0.05).

**Figure 2:**
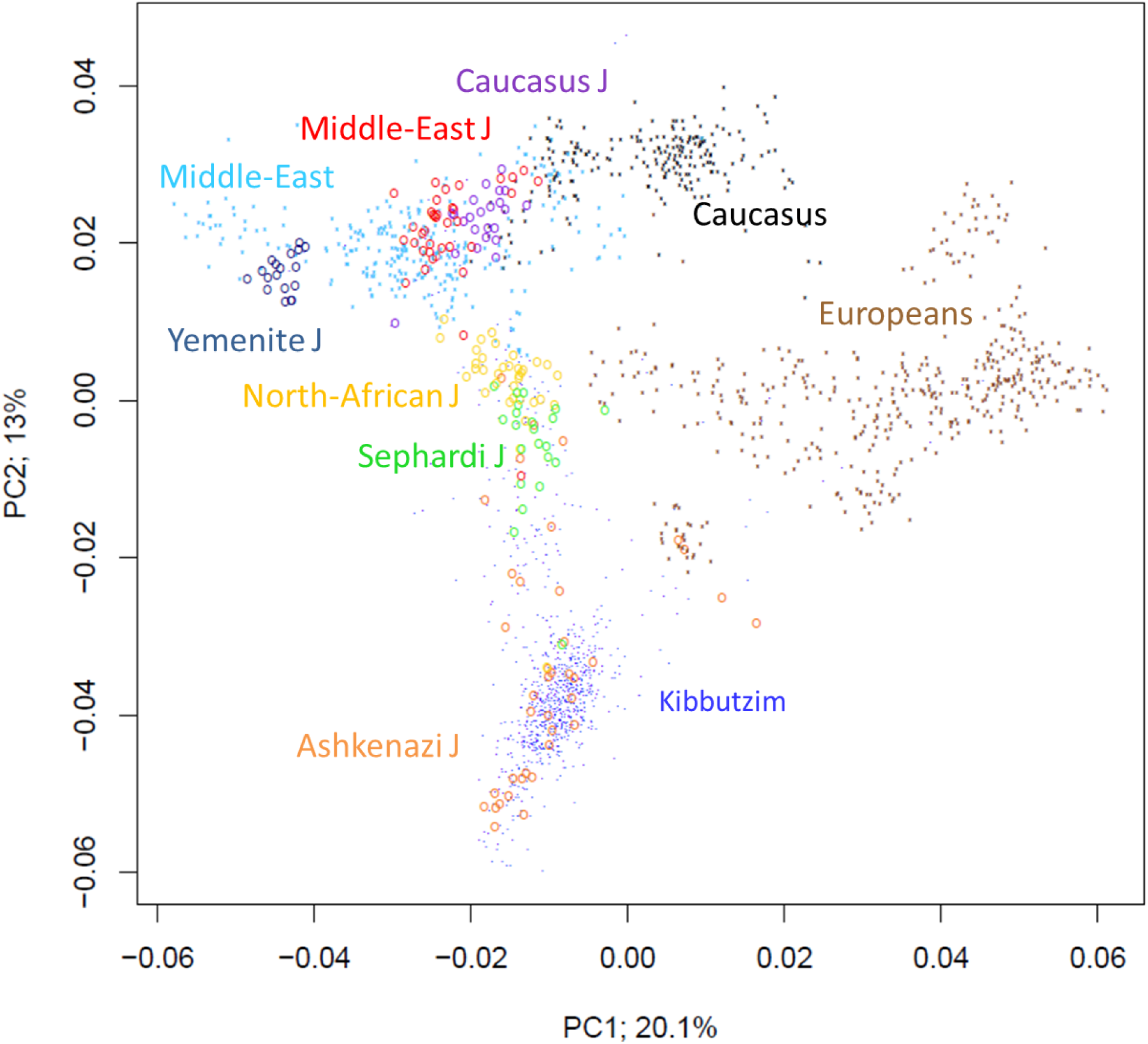
A plot of principal component 1 (PC1) and principal component 2 (PC2) for reference Jewish (n=174) and non-Jewish (n=922) populations, along with the KFS samples (n=901, blue dots).

**Figure 3:**
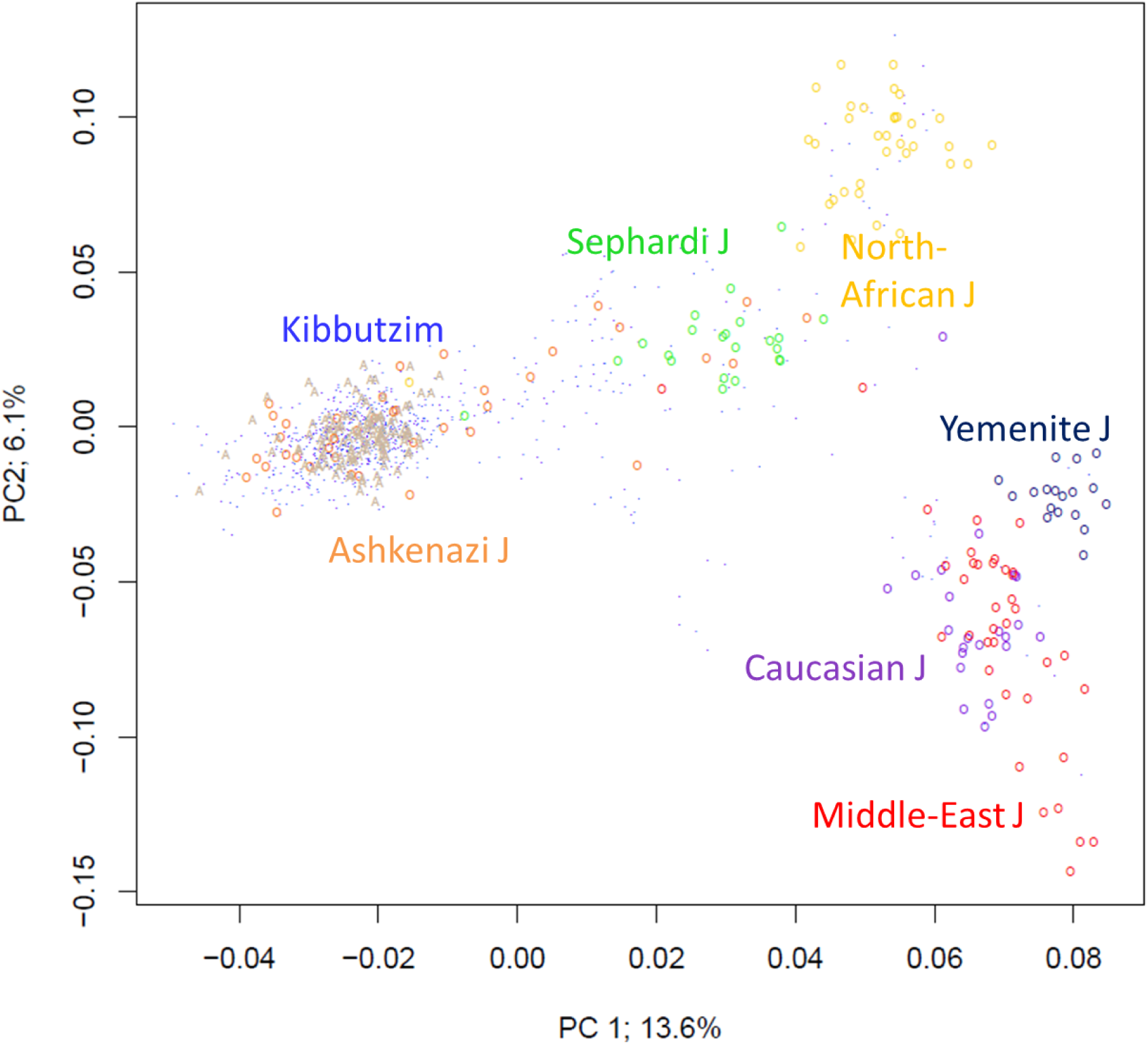
A PCA plot for the Jewish populations. Reference samples are as in Figure 2. Samples from The Ashkenazi Genome Consortium (TAGC) are shown in beige “A”s (n=128). The KFS samples are marked as blue dots (n=901), demonstrating that the ancestry of the KFS samples is predominantly (though not exclusively) Ashkenazi Jewish.

Next, we sought to determine whether the genetic ancestry of AJ with recent origins in Eastern Europe differs from those with recent origins in Western Europe. We designated KFS individuals born in Germany as Western AJ, and individuals born in Poland, Russia, Hungary, and Romania as Eastern AJ. A PCA plot revealed that Eastern and Western AJ can be distinguished in PC space, albeit imperfectly (Figure 4). Specifically, while many Western AJ clustered separately from Eastern AJ, some were indistinguishable from them. We observed the same pattern in the samples of (Behar *et al*. 2013).

**Figure 4:**
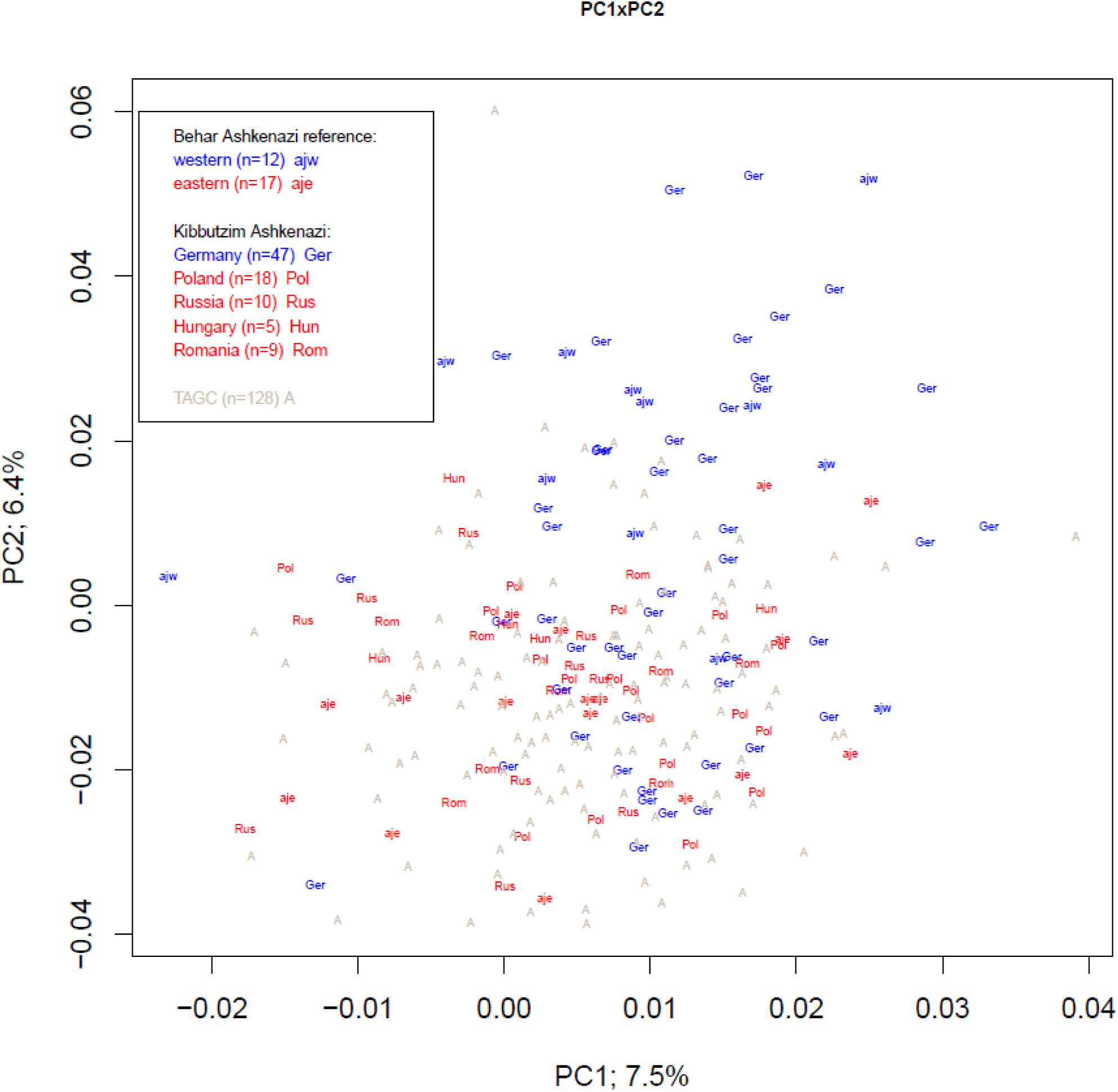
A PCA plot of the Ashkenazi Jewish individuals. The samples of Behar et al., 2013 are designated as “ajw” and “aje” for Western and Eastern Ashkenazi Jews, respectively. Samples from the KFS are indicated by their country of origin. For both datasets, Western and Eastern AJ samples are colored blue and red, respectively. Samples from The Ashkenazi Genome Consortium (TAGC) are marked with “A”s. The plot demonstrates that Western AJ have genetic ancestry slightly distinct from Eastern AJ, as many Western AJ cluster in the top-right quadrant. However, the distinction is imperfect.

We also sought to determine whether there are differences between the genetic ancestry of Israel-based and US-based AJ. To this end, we compared the (Israeli) KFS samples to a cohort of 128 US-based AJ who were recently whole-genome sequenced (Carmi *et al*. 2014). A PCA plot of the two groups, along with the Ashkenazi reference samples from Behar et al., 2013 (Figures 3 and 4), did not show any difference in the ancestry of Israel- and US-based AJ.

#### IBD Sharing and demographic reconstruction

To study sharing of identical-by-descent (IBD) haplotypes in the KFS study samples, we phased the genomes using SHAPEIT (with explicit modeling of parent-child relationships), and detected pairwise IBD segments longer than 3cM using Germline (Methods). The mean number of segments shared between a pair of unrelated KFS individuals (n=392 founders) was 4.19, with a mean segment length of 4.16cM.

It was previously shown that the number and lengths of IBD segments can be used to estimate the parameters of the founder event experienced by AJ (Palamara *et al*. 2012; Carmi *et al*. 2014). The time of the founder event was estimated as ≈25–35 generations ago, and the effective population size (the “bottleneck size”) as ≈300–400 individuals. We sought to determine whether these conclusions hold for our KFS AJ cohort, in particular that our study is family-based, which is expected to improve the accuracy of phasing and thereby IBD sharing detection (Methods).

Focusing on Ashkenazi founders (n=303; PC1 ≤ 0), we followed the method of (Palamara *et al*. 2012). We computed the proportion of the genome in IBD segments in each of a number of length intervals, and used the theory developed in (Palamara *et al*. 2012) and a grid search to find the best fitting demographic model (Supplementary Figure 2 and Methods). We limited ourselves to models with a constant ancestral size followed by a sudden reduction in the effective population size (a bottleneck) and an exponential expansion until the present (Supplementary Figure 3). The fitting errors for each model parameter are plotted in Supplementary Figure 4.

The inferred time of the founder event was 23 generations ago, which roughly corresponds to ≈600–700 years ago, slightly more recent than previous studies. The 95% confidence interval, using 100 bootstrap iterations over the chromosomes, was [22,24]. The inferred bottleneck size was 450 (diploid) individuals (bootstrap 95% confidence interval: [375,475]). This is slightly higher than in previous studies (≈1.5x), but still represents a severe bottleneck. The growth rate was estimated as 42% per generation (current effective size ≈1.7M, 95% interval: [0.4,2.6]M). The inferred ancestral population size was large (>10^5^, Supplementary Figure 4A), though we could not obtain a precise estimate, since the best fit was always on the upper boundary of the grid. This is likely due to the relatively little sharing of short IBD segments (≈3–4cM; Supplementary Figure 3) and the lack of information on even shorter segments (<3cM). Incorporating more complex models, shorter segments, and/or sequencing data is expected to improve the accuracy of the inferred parameters and resolve the population size history also between the bottleneck and ≈200 generations ago (Palamara *et al*. 2012; Browning and Browning 2015).

IBD sharing can also reveal differences in ancestry between Eastern and Western AJ. The mean number of segments shared within Western AJ was 1.4x larger than within Eastern AJ (8.4 vs 5.9, P<10^−7^; Table 2), but the mean segment length was similar (≈5.5cM, P=0.28). Sharing levels were particularly high in the group of Western AJ that was distinct in the PCA plot (Table 2). The number and lengths of runs of homozygosity (length >5Mbp; Methods) did not significantly differ between Eastern and Western AJ (2.18 vs. 2.21 segments with mean lengths 9.7 vs. 8.4Mbp, respectively; P>0.05 for both comparisons; Table 3).

**Table 2:** IBD sharing within Eastern and Western AJ. The mean number of segments and the mean segment length are shown for segments shared within each group. Sharing within Western AJ (WAJ) is also shown separately for “distinct” WAJ (defined as those having coordinates PC1>-0.01 and PC2>0.019 in Figure 4; n=11) and for WAJ similar to Eastern AJ (EAJ; all other WAJ; n=34). The The P-value for a difference between EAJ and (all) WAJ was calculated by permuting the population labels 10^7 times. The P-value was <10^-7 for comparing the number of segments and 0.28 for comparing the segment lengths.

|  | Number of segments | Segment length (cM) |
| --- | --- | --- |
| Eastern AJ | 5.87 | 5.50 |
| Western AJ | 8.39 | 5.59 |
| WAJ distinct | 12.40 | 5.71 |
| WAJ similar to EAJ | 7.43 | 5.57 |

**Table 3:** Runs of homozygosity in Western and Eastern AJ. The mean number of segments per individual and the mean segment length are shown for each group.

|  | Number of segments | Segment length (Mb) |
| --- | --- | --- |
| Eastern AJ | 2.21 | 8.368 |
| Western AJ | 2.18 | 9.769 |
| Wilcoxon test P-value | 0.15 | 0.29 |

### Functional Annotation of variants enriched in the KFS compared to Europeans

We annotated the function of 212,505 variants of minor allele frequency (MAF) > 1% in the KFS and with >10x MAF ratio between the KFS and Europeans (Methods). We identified 62 (0.03%) high impact and 291 (0.13%) moderate impact variants, with the remaining variants predicted to have low or no functional significance (“modifiers”) according to SnpEff (Supplementary Table 5). We observed no correlation between MAF enrichment in the KFS population and the putative functional significance (Supplementary Figure 5).

Gene-set enrichment analysis (GSEA) on 85 genes with >100x enriched variants and high/moderate functional impact (Methods) identified significant enrichment (false discovery rate q-value <0.01) within 32 gene sets (pathways) in the molecular signature database (MSigDB) in GSEA. The top pathways enriched for genes with AJ-specific variants were largely involved in cancers (breast cancer, pancreatic cancer, melanoma, and leukemia, among others). GSEA for >10x enriched variants with high or moderate functional significance returned similar pathways (not shown). There was no enrichment for cancer-related pathways when random sets of genes with variants of no functional significance were analyzed.

### Association of variants enriched in the KFS with anthropometric and cardiometabolic traits

We considered only the 212,505 variants enriched with >10x higher frequency in the KFS compared to Europeans, and used BOLT-LMM to test for an association with each of 16 anthropometric and cardiometabolic traits (Methods; qq-plots are shown in Supplementary Figure 6). Due to the relatively small number of variants tested, the P-value threshold for genome-wide significance was 1.61·10^−6^ (Methods). At this significance level, 24 variants demonstrated significant associations (Table 4).

**Table 4:**
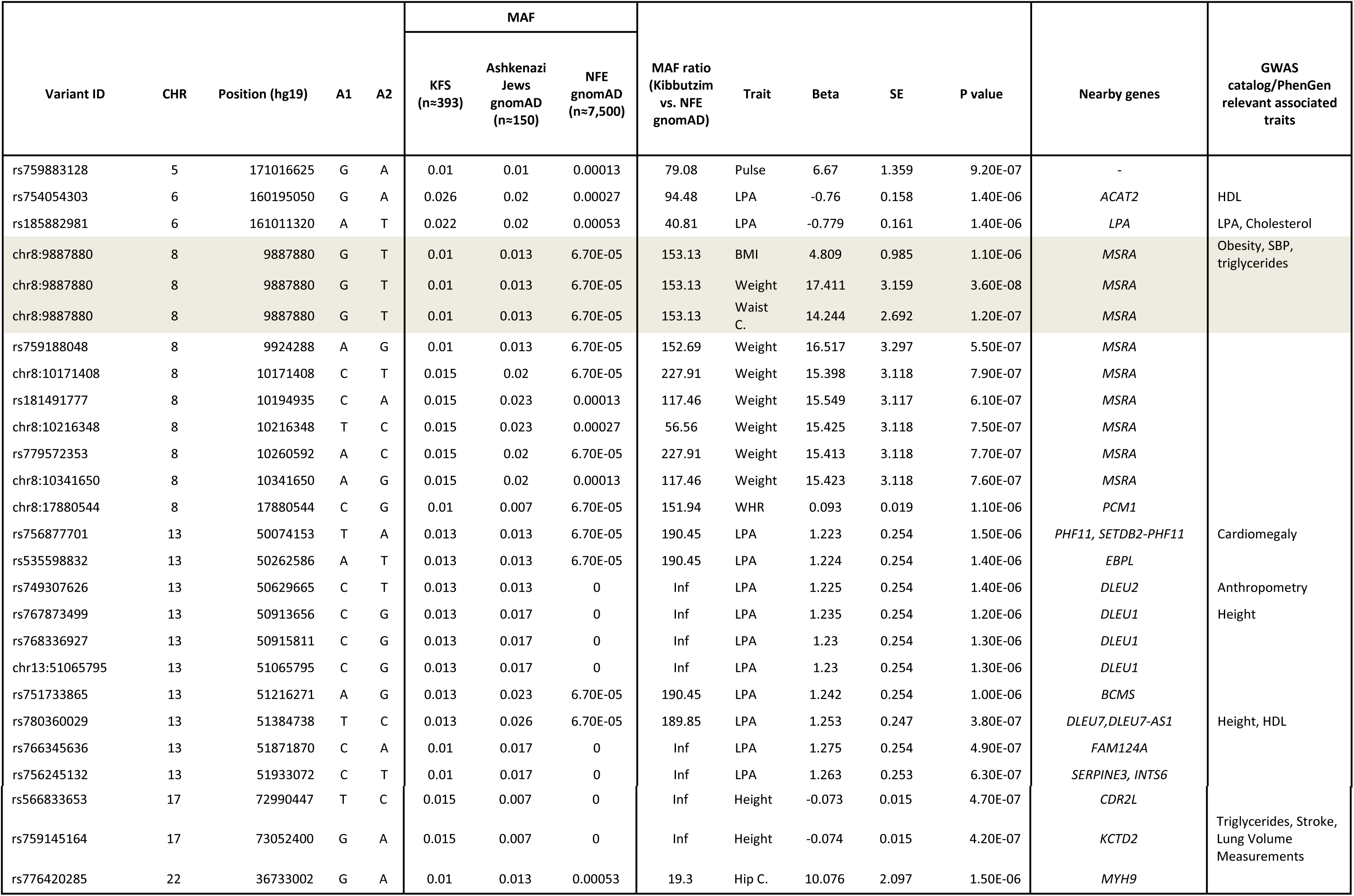
Genome-wide significant associations for enriched variants in the KFS (MAF > 1% and 10x higher compared to Europeans). A1 is the minor allele and the effect allele. Effect sizes are in units of the phenotypes (Supplementary Table 3). We used a genome-wide significance threshold of P=1.61·10^−6^ (Methods). The MAF in the KFS was computed in founders only. None of the variants was found in the 1000 Genomes NFE population. The most strongly associated variant, which is associated with BMI, weight, and waist circumference, is highlighted. Abbreviations: C.: circumference, WHR: waist to hip ratio, LPA: Lipoprotein(a)

Our main findings include a region spanning seven variants (453kb, MAF enrichment between 56 and 228), located in chr8p23.1, and associated with body weight, waist circumference, and body mass index (BMI). The most significant association was observed with body weight for (hg19) chr8:9887880:G>T (P = 3.6·10^−8^), an imputed variant located upstream of the *MSRA* (methionine sulfoxide reductase A) gene. This is the only variant with study-wide significant association (P<1.61·10^−6^/16), and it was also associated with waist circumference (P=1.2·10^−7^) and BMI (P=1.1·10^−6^). We replicated this finding in another Israeli cohort, the Jerusalem Perinatal Study (JPS), which we described in (Lawrence *et al*. 2015), and which consists mainly of parents and their children who were born in the 1970s in Jerusalem. Applying the same methodology as used here (a linear mixed model with adjusting for age and sex), chr8:9887880:G>T showed a nominally significant association in the JPS with weight (P=0.003, n=1291), BMI (P=0.035, n=1288), and waist circumference (P=0.038, n=448).

Another large region (1.9Mb) in chr13q14.3 showed a significant association with lipoprotein(a) (LPA). The region contained ten variants, all with >189-fold higher MAF between KFS and NFE, with the most significant result at rs780360029:C>T (P = 3.8·10^−7^). These variants span eight genes (Table 4), three of which belong to a region that is frequently deleted in B-cell chronic lymphocytic leukemia (called *DLEU*) (Rowntree *et al*. 2002). Two intronic variants in chr6q25.3-26 were associated with LPA; rs754054303 located in the *ACAT2* gene and rs185882981 located in the *LPA* gene (Table 4). Two additional intronic variants in chr17q25.1 — rs566833653 in *CDR2L* and rs759145164 in *KCTD2* — were both associated with height (P = 4.7·10^−7^ and P = 4.2·10^−7^, respectively) (Table 4), and are absent in Europeans.

## Discussion

We analyzed the genotypes of 901 individuals from extended families living in Kibbutzim in Israel, who participated in a longitudinal study with detailed records of anthropometric traits and cardiometabolic factors. The data enabled us to refine population genetic patterns of Israeli Jews, as well as study genetic associations with 16 traits.

### Ashkenazi Jewish population genetics

A PCA analysis confirmed self-reported ancestries (whenever available) and allowed precise assignment of ethnic origins for most individuals. Participants in this cohort were mostly of AJ origin (81.4%), with the remaining having various other Jewish ancestries. Present-day AJ descend from Jewish ancestors who lived in Eastern and (most of) Western Europe. One theory of Ashkenazi origins is an initial settlement in Western Europe (Northern France and Germany), and later migration and expansion in Poland and the rest of Eastern Europe (Weinryb 1972). An open question is whether the genetic ancestry of AJ with origins in Eastern Europe differs from those with recent origins in Western Europe. A previous study of ≈1300 AJ (Guha *et al*. 2012) did not find a correlation (on a PCA plot) between genetic ancestry and a country of origin. A later study of 29 AJ (Behar *et al*. 2013), which is part of the Jewish reference panel used here, did not identify genetic differences between Eastern and Western AJ, except for a minute East-Asian component in the ADMXITURE analysis that was present in Eastern, but not Western AJ. Our analysis of the KFS individuals who reported their country of origin showed that many Western AJ cluster separately from Eastern AJ, and the same pattern was observed in our re-analysis of the samples of (Behar *et al*. 2013). IBD sharing analysis showed, on average, 1.43x more shared segments within Western AJ compared to Eastern AJ, and an even higher levels of sharing (2.11x) for those Western AJ who were distinct on PCA (Table 2; Figure 4). These results suggest that at least some Western AJ descend from a slightly smaller subset of the founders compared to Eastern AJ (Table 2). An explanation consistent with these observations is that Western AJ consist of two slightly distinct groups: one that descends from a subset of the original founders (represented by those who are distinct on the PCA plot), and another that has migrated there back from Eastern Europe, possibly after receiving a limited degree of gene flow.

Following the large-scale migrations of the 20th century, present-day AJ live mostly in the US and in Israel. We compared the KFS samples, which were collected in Israel, to genomes of US-based AJ (Carmi *et al*. 2014). A PCA plot of the two groups (Figures 3 and 4) revealed no differences in the ancestry of Israel-based and US-based AJ. This result, which agrees with the IBD-based analysis of (Gusev *et al*. 2012), is expected based on the short time since the migrations out of Europe and suggests that the source population for these migrations was relatively homogeneous.

Sharing of IBD haplotypes between and within populations can be highly informative on recent demographic events (e.g., (Palamara *et al*. 2012; Ralph and Coop 2013; Browning and Browning 2015; Zidan *et al*. 2015)). It was previously estimated that AJ have experienced a founder event ≈25-35 generations ago with an effective population size ( “bottleneck size”) of ≈300-400 individuals (Palamara *et al*. 2012; Carmi *et al*. 2014). We established that these conclusions hold for our independent AJ sample, which is particularly important since our study is family-based, and thus has higher accuracy of IBD sharing detection.

### Analysis of enriched variants

Studying isolated and founder populations, such as the AJ population, is expected to increase power to discover disease-associated genes due to drift to relatively high frequencies of rare or unique risk alleles (Peltonen *et al*. 2000; Zeggini 2012; Hatzikotoulas *et al*. 2014). Additionally, linkage disequilibrium in isolated populations tends to extend over longer distances resulting in longer haplotypes, thus empowering imputation approaches. Here, we did not observe correlation between variants enriched in AJ (the KFS) and their putative functional significance. However, for functionally significant variants that are particularly enriched in AJ (>100x frequency ratio to Europeans), we identified a significant overlap with several cancer-related gene-sets (including breast cancer). AJ women have a much higher-than-average risk of breast cancer mostly due to a higher risk of having founder mutations in the *BRCA1* and *BRCA2* genes (Levy-Lahad *et al*. 1997). While no MAF enrichment was observed in *BRCA1* or *BRCA2* genes in KFS, some genes with enriched variants were found to interact with BRCA1 (Supplementary Table 6). We note that whether cancer is more prevalent in Ashkenazi Jews compared to the general Western population is debated, and possibly limited to colorectal and pancreatic cancers, if at all (Feldman 2001; Lynch *et al*. 2004; Streicher *et al*. 2017).

Overall, we observed 24 enriched variants in the KFS (compared to Europeans) that were associated with anthropometric and cardiometabolic traits. Regions of interest include seven variants located in chr8p23.1, surrounding the *MSRA* gene, which were associated with body weight, waist circumference, and BMI. The most significant variant, located in this region (chr8:9887880:G>T) was also replicated in another Israeli cohort, supporting our finding. Variants relatively close to this region (100kb upstream), located between the genes *TNKS* and *MSRA*, have been previously shown to be associated with extreme obesity in children and adolescents (Scherag *et al*. 2010) and with adult waist circumference (Lindgren *et al*. 2009) in European ancestry. Our findings may implicate *MSRA* as a candidate gene for the observed associations. This gene encodes a ubiquitous and highly conserved protein that carries out the enzymatic reduction of methionine sulfoxide to methionine. Another region of interest is chr13q14.3, showing significant associations of 10 variants with LPA. This region includes the *DLEU* genes, which are frequently deleted in B-cell chronic lymphocytic leukemia, suggesting a role of one or more tumor suppressors (Rowntree *et al*. 2002). The variant rs749307626 is located in an intronic region of the *DLEU2* gene, which was previously associated with waist-to-hip ratio in a meta-analysis of African and European populations (Ng *et al*. 2017). Three variants are located in the *DLEU1* gene, previously associated with anthropometric traits in another isolate; Korčula Island, Croatia (Polasek *et al*. 2009). One variant is located in the *DLEU7* gene, previously associated with height in Europeans and Africans (Kang *et al*. 2010). The variant rs756877701 is located in an intronic region of the *PHF11* gene, which was previously associated with cardiomegaly in the Amish population (Parsa *et al*. 2011). Two intronic variants in chr6q25.3-26 that were associated with LPA are located in the known *LPA* locus (Ober *et al*. 2009; Kettunen *et al*. 2016), providing evidence for the generalizability of our results. Other variants (non-enriched) in this region also showed highly significant associations with LPA in the KFS; the most significant association was seen for rs56393506:C>T, 1900bp upstream of *LPA’s* 5’UTR, P= 9.1·10^−17^.

### Future directions

We reported the first genome-wide association analysis (to our knowledge) of cardiometabolic traits in the Israeli Jewish population, and detected a number of suggestive associations. Our study also refined the understanding of the population genetics of Ashkenazi and other Jewish groups. Current limitations of our study include its relatively small sample size and its focus on Ashkenazi Jews. Thus, additional analyses will be required in larger Jewish samples, as well as in other populations, to replicate the findings and elucidate the role of the associated variants in the development of the phenotype.

We envision a number of ways in which our data could be used in future studies by researchers interested in the genetics of anthropometric and cardiometabolic traits, as well as in Jewish genetics. First, our cohort could be meta-analyzed along with larger cohorts, or could be used for the replication of signals found in other populations. Power to replicate associations in our cohort is expected to be relatively high, due to the Ashkenazi founder event (Carmi *et al*. 2014) and the relatively homogenous environment of the Kibbutzim. Second, our data could be used to improve the understanding of differences in the genetic architecture of the investigated traits between Jews and other populations. Similarly, our data could be used to compare and refine the genomic prediction accuracy across populations (Marquez-Luna *et al*. 2016; Coram *et al*. 2017). Third, the family structure of our cohort and the presence of longitudinal phenotypic data will allow additional analyses, such as a GWAS on the longitudinal change and a search for parent of origin effects. Finally, our cohort could be utilized for studies of the population genetics of the Jewish people. While other cohorts exist (Atzmon *et al*. 2010; Guha *et al*. 2012; Behar *et al*. 2013), our cohort is the largest to include extended families, it represents multiple Jewish ancestries, and it has (partial) information on the participants’ countries of origin. For example, algorithms for identification of related individuals in a random sample are known to underperform in AJ, due to the abundance of IBD sharing even between unrelated individuals (Paull *et al*. 2014). Our dataset could be used for the development of algorithms for pedigree reconstruction in AJ and other isolated populations.

## Acknowledgements

We are grateful to the study participants, recruiters, interviewers and nurses. This study was supported by Israeli Science Foundation grant 201/98-1 and partially by National Institutes of Health research grant R01HL088884. Genotyping was also supported in part by a generous gift from the Samson Family (South Africa) to DK.

## Additional information

### Conflict of interest

None declared.

### Data availability

We are in the process of submitting the data reported in this paper to the European Genome-phenome Archive (EGA).

## References

Atanasovska B., Kumar V., Fu J., Wijmenga C., Hofker M. H., 2015 GWAS as a Driver of Gene Discovery in Cardiometabolic Diseases. Trends Endocrinol. Metab. 26: 722–732.

Atzmon G., Hao L., Pe’er I., Velez C., Pearlman A., et al., 2010 Abraham’s Children in the Genome Era: Major Jewish Diaspora Populations Comprise Distinct Genetic Clusters with Shared Middle Eastern Ancestry. Am. J. Hum. Genet. 86: 850–859.

Auton A., Abecasis G. R., Altshuler D. M., Durbin R. M., Abecasis G. R., et al., 2015 A global reference for human genetic variation. Nature 526: 68–74.

Baier L. J., L. Hanson R., 2004 Genetic Studies of the Etiology of Type 2 Diabetes in Pima Indians Hunting for Pieces to a Complicated Puzzle. Diabetes 53: 1181–1186.

Behar D. M., Metspalu M., Baran Y., Kopelman N. M., Yunusbayev B., et al., 2013 No evidence from genome-wide data of a Khazar origin for the Ashkenazi Jews. Hum. Biol. 85: 859–900.

Browning S. R., Browning B. L., 2015 Accurate Non-parametric Estimation of Recent Effective Population Size from Segments of Identity by Descent. Am. J. Hum. Genet. 97: 404–418.

Carmi S., Hui K. Y., Kochav E., Liu X., Xue J., et al., 2014 Sequencing an Ashkenazi reference panel supports population-targeted personal genomics and illuminates Jewish and European origins. Nat. Commun. 5: 4835.

Chang C. C., Chow C. C., Tellier L. C., Vattikuti S., Purcell S. M., et al., 2015 Second-generation PLINK: rising to the challenge of larger and richer datasets. Gigascience 4: 7.

Charrow J., 2004 Ashkenazi Jewish genetic disorders. Fam. Cancer 3: 201–206.

Cingolani P., Platts A., Wang L. L., Coon M., Nguyen T., et al., 2012 A program for annotating and predicting the effects of single nucleotide polymorphisms, SnpEff: SNPs in the genome of Drosophila melanogaster strain w1118; iso-2; iso-3. Fly (Austin).

Conomos M. P., Reiner A. P., Weir B. S., Thornton T. A., 2016 Model-free Estimation of Recent Genetic Relatedness. Am. J. Hum. Genet. 98: 127–148.

Coram M. A., Fang H., Candille S. I., Assimes T. L., Tang H., 2017 Leveraging Multi-ethnic Evidence for Risk Assessment of Quantitative Traits in Minority Populations. Am. J. Hum. Genet. 101: 218–226.

Cui J. S., Hopper J. L., Harrap S. B., 2003 Antihypertensive treatments obscure familial contributions to blood pressure variation. Hypertension 41: 207–210.

Durand E. Y., Eriksson N., Mclean C. Y., 2014 Reducing pervasive false-positive identical-by-descent segments detected by large-scale pedigree analysis. Mol. Biol. Evol. 31: 2212–2222.

Fall T., Ingelsson E., 2014 Genome-wide association studies of obesity and metabolic syndrome. Mol. Cell. Endocrinol. 382: 740–757.

Fang S., Zhang S., Sha Q., 2016 Literature Reviews on Methods for Rare Variant Association Studies. Hum. Genet. Embryol. 6: 1–5.

Feldman G. E., 2001 Do Ashkenazi Jews have a higher than expected cancer burden? Implications for cancer control prioritization efforts. Isr. Med. Assoc. J. 3: 341–346.

Friedlander Y., Elkana Y., Sinnreich R., Kark J. D., 1995 Genetic and environmental sources of fibrinogen variability in Israeli families: the Kibbutzim Family Study. Am. J. Hum. Genet. 56: 1194–206.

Friedlander Y., Kark J. D., Sinnreich R., Edwards K. L., Austin M. a, 1999a Inheritance of LDL peak particle diameter: results from a segregation analysis in Israeli families. Genet. Epidemiol. 16: 382–96.

Friedlander Y., Lapidos T., Sinnreich R., Kark J. D., 1999b Genetic and environmental sources of QT interval variability in Israeli families: the kibbutz settlements family study. Clin. Genet. 56: 200–9.

Friedlander Y., Kark J. D., Sinnreich R., Basso F., Humphries S. E., 2003 Combined segregation and linkage analysis of fibrinogen variability in Israeli families: evidence for two quantitative-trait loci, one of which is linked to a functional variant (-58G > A) in the promoter of the alpha-fibrinogen gene. Ann. Hum. Genet. 67: 228–41.

Friedlander Y., Vatta M., Sotoodehnia N., Sinnreich R., Li H., et al., 2005 Possible association of the human KCNE1 (minK) gene and QT interval in healthy subjects: evidence from association and linkage analyses in Israeli families. Ann. Hum. Genet. 69: 645–56.

Friedlander Y., Kark J. D., Sinnreich R., Tracy R. P., Siscovick D. S., 2006 Fibrinogen and CRP in Israeli families: genetic and environmental sources of concentrations and longitudinal changes. Atherosclerosis 189: 169–77.

Gilbert E., Carmi S., Ennis S., Wilson J. F., Cavalleri G. L., 2017 Genomic insights into the population structure and history of the Irish Travellers. Sci. Rep. 7: 42187.

Gilly A., Ritchie G. R., Southam L., Farmaki A. E., Tsafantakis E., et al., 2016 Very low-depth sequencing in a founder population identifies a cardioprotective APOC3 signal missed by genome-wide imputation. Hum. Mol. Genet. 25: 2360–2365.

Guha S., Rosenfeld J. a, Malhotra A. K., Lee A. T., Gregersen P. K., et al., 2012 Implications for health and disease in the genetic signature of the Ashkenazi Jewish population. Genome Biol. 13: R2.

Gusev A., Lowe J. K., Stoffel M., Daly M. J., Altshuler D., et al., 2009 Whole population, genome-wide mapping of hidden relatedness. Genome Res. 19: 318–26.

Gusev A., Palamara P. F., Aponte G., Zhuang Z., Darvasi A., et al., 2012 The architecture of long-range haplotypes shared within and across populations. Mol. Biol. Evol. 29: 473–486.

Hatzikotoulas K., Gilly A., Zeggini E., 2014 Using population isolates in genetic association studies. Brief. Funct. Genomics 13: 371–377.

Howie B., Fuchsberger C., Stephens M., Marchini J., Abecasis G. R., 2012 Fast and accurate genotype imputation in genome-wide association studies through pre-phasing. Nat. Genet. 44: 955–959.

Kang S. J., Chiang C. W. K., Palmer C. D., Tayo B. O., Lettre G., et al., 2010 Genome-wide association of anthropometric traits in African- and African-derived populations. Hum. Mol. Genet. 19: 2725–2738.

Kark J. D., Carmel S., Sinnreich R., Goldberger N., Friedlander Y., 1996 Psychosocial factors among members of religious and secular kibbutzim. Isr. J. Med. Sci. 32: 185–94.

Kenny E. E., Pe’er I., Karban A., Ozelius L., Mitchell A. A., et al., 2012 A Genome-Wide Scan of Ashkenazi Jewish Crohn’s Disease Suggests Novel Susceptibility Loci. PLoS Genet. 8: 1–10.

Kettunen J., Demirkan A., Würtz P., Draisma H. H. M. M., Haller T., et al., 2016 Genome-wide study for circulating metabolites identifies 62 loci and reveals novel systemic effects of LPA. Nat. Commun. 7: 11122.

Kristiansson K., Naukkarinen J., Peltonen L., 2008 Isolated populations and complex disease gene identification. Genome Biol. 9: 109.

Lawrence G. M., Siscovick D. S., Calderon-Margalit R., Enquobahrie D. A., Granot-Hershkovitz E., et al., 2015 Cohort Profile: The Jerusalem Perinatal Family Follow-Up Study. Int. J. Epidemiol.: dyv120.

Lek M., Karczewski K. J., Minikel E. V., Samocha K. E., Banks E., et al., 2016 Analysis of protein-coding genetic variation in 60,706 humans. Nature 536: 285–291.

Lemaitre R. N., Siscovick D. S., Berry E. M., Kark J. D., Friedlander Y., 2008 Familial aggregation of red blood cell membrane fatty acid composition: the Kibbutzim Family Study. Metabolism. 57: 662–8.

Levy-Lahad E., Catane R., Eisenberg S., Kaufman B., Hornreich G., et al., 1997 Founder BRCA1 and BRCA2 mutations in Ashkenazi Jews in Israel: frequency and differential penetrance in ovarian cancer and in breast-ovarian cancer families. Am. J. Hum. Genet. 60: 1059–1067.

Lindgren C. M., Heid I. M., Randall J. C., Lamina C., Steinthorsdottir V., et al., 2009 Genome-wide association scan meta-analysis identifies three loci influencing adiposity and fat distribution. PLoS Genet. 5.

Loh P.-R., Bhatia G., Gusev A., Finucane H. K., Bulik-Sullivan B. K., et al., 2015a Contrasting genetic architectures of schizophrenia and other complex diseases using fast variance-components analysis. Nat. Genet. 47: 1385–92.

Loh P.-R., Tucker G., Bulik-Sullivan B. K., Vilhjálmsson B. J., Finucane H. K., et al., 2015b Efficient Bayesian mixed-model analysis increases association power in large cohorts. Nat. Genet. 47: 284–290.

Lopes F. L., Hou L., Boldt A. B. W., Kassem L., Alves M., et al., 2016 Finding Rare, Disease-Associated Variants in Isolated Groups: Potential Advantages of Mennonite Populations. Hum. Biol. 88: 109–120.

Lynch H. T., Rubinstein W. S., Locker G. Y., 2004 Cancer in Jews: introduction and overview. Fam. Cancer 3: 177–192.

Marquez-Luna C., Price A. L., Price A. L., 2016 Multi-ethnic polygenic risk scores improve risk prediction in diverse populations. bioRxiv: 51458.

Mathers C. D., Loncar D., 2006 Projections of Global Mortality and Burden of Disease from 2002 to 2030. Plos Med. 3: 2011–2030.

Nair A. K., Baier L. J., 2015 Complex genetics of type 2 diabetes and effect size: What have we learned from isolated populations? Rev. Diabet. Stud. 12: 299–319.

Ng M. C. Y., Graff M., Lu Y., Justice A. E., Mudgal P., et al., 2017 Discovery and fine-mapping of adiposity loci using high density imputation of genome-wide association studies in individuals of African ancestry: African Ancestry Anthropometry Genetics Consortium (GP Copenhaver, Ed.). PLOS Genet. 13: e1006719.

O’Connell J., Gurdasani D., Delaneau O., Pirastu N., Ulivi S., et al., 2014 A General Approach for Haplotype Phasing across the Full Spectrum of Relatedness. PLoS Genet. 10.

Ober C., Nord A. S., Thompson E. E., Pan L., Tan Z., et al., 2009 Genome-wide association study of plasma lipoprotein(a) levels identifies multiple genes on chromosome 6q. J. Lipid Res. 50: 798–806.

Ott J., Kamatani Y., Lathrop M., 2011 Family-based designs for genome-wide association studies. Nat. Rev. Genet. 12: 465–474.

Palamara P. F., Lencz T., Darvasi A., Pe’er I., 2012 Length distributions of identity by descent reveal fine-scale demographic history. Am. J. Hum. Genet. 91: 809–822.

Parsa A., Chang Y. P. C., Kelly R. J., Corretti M. C., Ryan K. A., et al., 2011 Hypertrophy-associated polymorphisms ascertained in a founder cohort applied to heart failure risk and mortality. Clin. Transl. Sci. 4: 17–23.

Paull J. M., Tannenbaum G., Briskman J., 2014 Differences in Autosomal DNA Characteristics between Jewish and Non-Jewish Populations Excerpted from: ‘Why Autosomal DNA Test Results are Significantly Different for Ashkenazi. SurnameDNA J. XXX: 1–32.

Pe’er I., Yelensky R., Altshuler D., Daly M. J., 2008 Estimation of the multiple testing burden for genomewide association studies of nearly all common variants. Genet. Epidemiol. 32: 381–385.

Peltonen L., Palotie a, Lange K., 2000 Use of population isolates for mapping complex traits. Nat. Rev. Genet. 1: 182–90.

Polasek O., Marusić A., Rotim K., Hayward C., Vitart V., et al., 2009 Genome-wide association study of anthropometric traits in Korcula Island, Croatia. Croat. Med. J. 50: 7–16.

Ralph P., Coop G., 2013 The Geography of Recent Genetic Ancestry across Europe. PLoS Biol. 11.

Rowntree C., Duke V., Panayiotidis P., Kotsi P., Palmisano G., et al., 2002 Deletion analysis of chromosome 13q14.3 and characterisation of an alternative splice form of LEU1 in B cell chronic lymphocytic leukemia. Leukemia 16: 1267–1275.

Sabatti C., Service S. K., Hartikainen A.-L., Pouta A., Ripatti S., et al., 2009 Genome-wide association analysis of metabolic traits in a birth cohort from a founder population. Nat. Genet. 41: 35–46.

Scherag A., Dina C., Hinney A., Vatin V., Scherag S., et al., 2010 Two new loci for body-weight regulation identified in a joint analysis of genome-wide association studies for early-onset extreme obesity in French and German study groups. PLoS Genet. 6: 2–11.

Sinnreich R., Friedlander Y., Sapoznikov D., Kark J. D., 1998 Familial aggregation of heart rate variability based on short recordings--the kibbutzim family study. Hum. Genet. 103: 34–40.

Sinnreich R., Friedlander Y., Luria M. H., Sapoznikov D., Kark J. D., 1999 Inheritance of heart rate variability: the kibbutzim family study. Hum. Genet. 105: 654–61.

Streicher S. A., Klein A. P., Olson S. H., Amundadottir L. T., DeWan A. T., et al., 2017 Impact of Sixteen Established Pancreatic Cancer Susceptibility Loci in American Jews. Cancer Epidemiol. biomarkers Prev. 10: 1540–1548.

Subramanian A., Tamayo P., Mootha V. K., Mukherjee S., Ebert B. L., et al., 2005 Gene set enrichment analysis: A knowledge-based approach for interpreting genome-wide expression profiles. PNAS 102: 15545–15550.

Trecartin R. F., Liebhaber S. A., Chang J. C., 1981 β° Thalassemia in Sardinia is caused by a nonsense mutation. J. Clin. Invest. 68: 1012–1017.

Weinryb B. D. (Bernard D., 1972 The Jews of Poland; a social and economic history of the Jewish community in Poland from 1100 to 1800. Jewish Publication Society of America.

Zeggini E., 2012 Europe PMC Funders Group Next-generation association studies for complex traits. Nat. Genet. 43: 287–288.

Zeggini E., 2014 Using genetically isolated populations to understand the genomic basis of disease. Genome Med. 6: 83.

Zeggini E., Gloyn A. L., Hansen T., 2016 Insights into metabolic disease from studying genetics in isolated populations: stories from Greece to Greenland. Diabetologia 59: 938–941.

Zidan J., Ben-Avraham D., Carmi S., Maray T., Friedman E., et al., 2015 Genotyping of geographically diverse Druze trios reveals substructure and a recent bottleneck. Eur. J. Hum. Genet. 23: 1–7.

